# Janus effect of glucocorticoids on differentiation of muscle fibro/adipogenic progenitors

**DOI:** 10.1101/581363

**Authors:** Andrea Cerquone Perpetuini, Alessio Reggio, Mauro Cerretani, Giulio Giuliani, Marisabella Santoriello, Roberta Stefanelli, Alessandro Palma, Steven Harper, Luisa Castagnoli, Alberto Bresciani, Gianni Cesareni

## Abstract

Muscle resident fibro-adipogenic progenitors (FAPs), support muscle regeneration by releasing cytokines that stimulate the differentiation of myogenic stem cells. However, in non-physiological contexts (myopathies, atrophy, aging) FAPs cause fibrotic and fat infiltrations that impair muscle function. We set out to perform a fluorescence microscopy-based screening to identify compounds that perturb the differentiation trajectories of these multipotent stem cells. From a primary screen of 1120 FDA/EMA approved drugs, we identified 34 compounds as potential inhibitors of adipogenic differentiation of FAPs isolated from the murine model (mdx) of Duchenne muscular dystrophy (DMD). The hit list from this screen was surprisingly enriched with compounds from the glucocorticoid (GCs) chemical class, drugs that are known to promote adipogenesis in vitro and in vivo. To shed light on these data, three GCs identified in our screening efforts were characterized by different approaches. We found that like dexamethasone, budesonide inhibits adipogenesis induced by insulin in subconfluent FAPs. However, both drugs have a proadipogenic impact when the adipogenic mix contains factors that increase the concentration of cAMP. Gene expression analysis demonstrated that treatment with glucocorticoids induces the transcription of Gilz/Tsc22d3, an inhibitor of the adipogenic master regulator PPARγ, only in anti-adipogenic conditions. Additionally, alongside their anti-adipogenic effect, GCs are shown to promote terminal differentiation of satellite cells. Both the anti-adipogenic and pro-myogenic effects are mediated by the glucocorticoid receptor and are not observed in the presence of receptor inhibitors. Steroid administration currently represents the standard treatment for DMD patients, the rationale being based on their anti-inflammatory effects. The findings presented here offer new insights on additional glucocorticoid effects on muscle stem cells that may affect muscle homeostasis and physiology.

## Introduction

In muscular dystrophies the degeneration of muscle tissue is initially compensated by efficient regeneration that neutralize muscle loss ^1^. However, over time, the regenerative potential in muscles of patients affected by myopathies is impaired and myofibers repair is curbed by the formation of fibrotic scars and fat infiltrations, ultimately leading to decreased muscle function ^2^. Muscle fibro-adipogenic progenitors play an important role in these processes. FAPs are muscle mesenchymal stem cells residing in the muscle fibers interstitial space. FAPs express the SCA1, CD34 and PDGFRα (CD140a) antigens while they are negative for the hematopoietic and endothelial markers, CD45 and CD31, and for the satellite marker α7 integrin (ITGA7) ^2–4^. FAPs contribute to muscle regeneration by secreting IGF-1 and IL-6 ^3^, by facilitating the clearance of necrotic debris ^5^ and by promoting the formation of extra-cellular matrix ^6^. In addition to this pro-regenerative roles, FAPs are responsible for the formation of ectopic tissue infiltrations in degenerating dystrophic muscles ^6^. For these reasons, drugs targeting the FAPs fibro-adipogenic potential are considered in clinical trials to alleviate the degeneration of muscle function in dystrophic patients ^7^. Recent studies have linked histone deacetylase inhibitors (HDACi) to a complex epigenetic network that modulates FAPs fibro-adipogenic differentiation in muscular dystrophies ^8–10^. In particular, the HDACi Trichostatin A (TSA) promotes the expression of two components of the myogenic transcriptional machinery, MyoD and BAF60C, a subunit of the SWI/SNF chromatin-remodeling complex promoting the switch from a fibro-adipogenic to a pro-myogenic phenotype ^10–12^. To alleviate the unfavorable consequences of fat infiltrations in myopathies it would be desirable to enrich our toolbox of drugs controlling FAPs adipogenesis by alternative mechanisms. To this end we performed a screening looking for new modulators of the fibro-adipogenic differentiation of FAPs isolated from the *mdx* mice model of Duchenne muscular dystrophy. Much to our surprise we observed an enrichment in glucocorticoids among the molecules observed to be negative modulators of adipogenic differentiation. Since glucocorticoids are often described as promoters of adipogenesis, we set out to shed light on this “Janus-like effect” of glucocorticoids on the differentiation of adipocyte progenitors. Steroids presently represent the standard pharmacological treatment for DMD patients ^13^. Despite their moderate beneficial effect on disease progression, their etiological role is not well understood. Here we present results suggesting that FAPs are a glucocorticoid target and that the anti-adipogenic effect of this class of molecules may contribute to their beneficial impact in delaying DMD progression.

## Materials and methods

### Mouse strains

In all the experiments young (6 weeks old) C57BL/6ScSn-Dmd^mdx^/J (*mdx* mice) or C57BL/6 (wt mice) purchased from the Jackson Laboratories were used. Mice were maintained according to standard animal facility procedures and experiments on animals were conducted according to the rules of good animal experimentation I.A.C.U.C. n°432 of March 12 2006. All experimental protocols were approved by the internal Animal Research Ethical Committee according to the Italian Ministry of Health regulation.

### Satellite cell and FAPs isolation

Hind limb muscles were isolated from *mdx* or wt mice. Muscles were then subjected to mechanical dissociation followed by enzymatic digestion for 60 minutes at 37°C. The enzymatic mix was composed by 2 μg/ml collagenase A (Roche 10103586001), 2,4 U/ml dispase II (Roche 04942078001) and 0,01 mg/ml DNase I (Roche 04716728001) in D-PBS with Calcium and Magnesium (Biowest L0625-500). Enzymatic digestion was stopped by addition of Hank’s Balanced Salt Solution (Thermo Fisher Scientific 14025050) and cell suspension was filtered through a 100, 70, 40 μm pores cell strainer. Red Blood Cells were removed using RBC Lysis Buffer (Santa Cruz sc-296258) and cell suspension was filtered through 30 μm pore cell strainer. Cells were sorted using the MACS separation technology. The antibodies used were CD45 (Miltenyi 130-052-301), CD31 (Miltenyi 130-097-418), ITGA7 (Miltenyi 130-104-261) and Sca1 (Miltenyi 130-106-641). Satellite cells (SCs) were purified as CD45−/CD31−/ITGA7+ cells, while FAPs were selected as CD45−/CD31−/ITGA7−/Sca1+ cells.

### Cell culture

For *in vitro* expansion, freshly isolated *mdx* FAPs (P0) were plated at a density of 2*10^5^ cells in a 10 cm Petri dish and cultured for four additional days in Cytogrow (Resnova TGM-9001-B). Cells were detached (P1) and used for specific experimental procedures.

To induce adipogenic differentiation, *mdx* FAPs were cultured at 37°C and 5% CO_2_, in growth medium (fGM) containing Dulbecco modified Eagle medium (DMEM) supplemented with 20% heat-inactivated fetal bovine serum (FBS) (Euroclone, #ECS0180L), 100 U/ml penicillin, 100 mg/ml streptomycin, 1mM sodium pyruvate and 10mM HEPES and 1 μg/mL insulin (Sigma, #I9278). Alternatively, FAPs were cultured in DMEM with 20% FBS, without insulin (GM), for 6 days. Confluent cells were then exposed to an Adipogenic Induction Medium (AIM) consisting of 10% FBS, 1 μg/mL insulin and 0.5 mM of 3-isobutyl-1-methylxanthine (IBMX) (Sigma, #I5879) complemented with budesonide (Selleck Chemicals, #S1286) or dexamethasone (Sigma, #D4902), for two days. After 48 hours, cells were switched to maintenance medium (MM) consisting of DMEM with 10% FBS and 1 μg/mL insulin for further 48 hours.

Satellite cells were cultured at 37°C and 5% CO_2_ on matrigel-coated plates in Satellite cell Growth Medium (sGM) composed of DMEM, 20% FBS, 10% horse serum (Euroclone, #ECS0090D), 1% Chicken Embryo Extract (Seralab, # CE-650-J). Prior to starting any experiment, freshly isolated SCs were cultured for at least 48 hours before treatment to allow cell adhesion.

The C2C12 mouse myoblast cell line was purchased from ATCC (American Type Culture Collection, Bethesda, MD, USA) company (CRL-1772). C2C12 were seeded on Falcon dishes at 37°C with 5% CO_2_ in growth medium (cGM) composed of DMEM supplemented with 10% FBS, 100 U/ml penicillin, 100 mg/ml streptomycin, 1mM sodium pyruvate and 10mM HEPES. C2C12 or SCs were differentiated into myotubes by growing them in cGM or sGM, respectively and let to differentiate spontaneously.

3T3-L1 cell line was obtained from the American Type Culture Collection (ATCC, CL-173™) and cultured at 37 °C in 5% CO_2_ atmosphere using pre-adipocyte expansion medium consisting of DMEM supplemented with 10% bovine calf serum, 100 U/ml penicillin and 100 mg/ml streptomycin. Pre-adipocytes were induced to differentiate following the protocol provided by ATCC. Pre-adipocytes were growth until 100% confluent and fed with pre-adipocyte expansion medium for further 48 hours. Pre-adipocytes were then incubated for 48 hours with differentiation medium consisting of: DMEM, 10% FBS, 1.0 μM dexamethasone, 0.5 mM IBMX and 1.0 μg/mL of insulin.

### Prestwick screening and drug treatment

To screen the Prestwick library *mdx* FAPs were seeded on 384 well plate at the density of 1500 cells/well. 24 hours after seeding cells were treated for 6 additional days with the 1120 compounds of the Prestwick library at the concentration of 5 μM. DMSO and TSA have been used as negative and positive controls respectively. Compounds were transferred from a 10 mM DMSO stock solution to assay plates by acoustic droplet ejection (ATS-100, EDC biosystems, USA). Cells were stained with ORO to visualize adipocytes while Hoechst 33342 was used to stain nuclei. Adipogenic differentiation has been assessed as the total pixel intensity (TPI) for ORO signal normalized for the number of nuclei in each field (ORO^norm^). Adipogenic differentiation has been reported in Table S1 as:

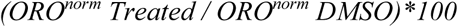

For validation studies, cells were treated with budesonide dissolved in DMSO, dexamethasone dissolved in ethanol, or mifepristone (RU-486, Selleck Chemicals, #S2606) dissolved in DMSO, at various concentrations and administered at specific times, as indicated in the figure legends.

### Immunoblotting

After the removal of culture medium, cells were washed in plate with PBS and homogenized in lysis buffer (Millipore cell signaling lysis buffer, #43-040) or RIPA buffer supplemented with protease inhibitor cocktail 200X (Sigma, #P8340) and phosphatase inhibitor cocktail I and II 100X (Sigma, #P5726, #P0044). Samples were then incubated in ice for 30 minutes with the lysis buffer and the cell debris separated by centrifugation at 17968 x g for 30 minutes, at 4°C. Protein concentrations were determined by Bradford colorimetric assay (Bio-Rad, #5000006). Total protein extracts (15μg or 20μg) were then separated by SDS-PAGE. Gels were transferred to nitrocellulose membranes saturated with blocking solution (5% milk or BSA and 0,1% Tween-20 in PBS) and incubated with primary antibodies overnight at 4°C. The antibodies used were as follows: mouse anti-vinculin (1:1000, Abcam, #ab18058), rabbit anti-perilipin (1:1000, Cell Signaling, #3470), mouse anti-Smooth Muscle Actin (SMA) (1:1000, Sigma, #A5228) and rat anti-Gilz (1:250, Invitrogen, #14-4033-80). Following the incubations with primary antibodies, membranes were then washed three times with the washing solution (0,1% Tween-20 in PBS) and incubated with anti-mouse or anti-rabbit secondary antibodies conjugated with HRP (horseradish peroxidase) (1:2500, Jackson ImmunoResearch) or anti-rat secondary antibody conjugated with HRP (1:10000, Invitrogen, #18-4818-82) for 1h at RT. The blots were further washed three times and visualized with an enhanced chemiluminescent immunoblotting detection system (Bio-Rad, #1705061). Densitometric analysis was performed using ImageJ software. Vinculin was used as a normalization control.

### Immunofluorescence

Cells were fixed with 2% paraformaldehyde (PFA) for 10 minutes at Room Temperature (RT) and permeabilized in 0,1% Triton X-100 for 5 minutes. Samples were then blocked with PBS, 10% FBS 0,1% TritonX-100 for 1 h at RT. Incubation with the primary antibody was performed for 1 h at RT, then cells were washed three times and incubated with the secondary antibody for 30 minutes at RT. The antibodies used were the following: mouse anti-myogenin (1:250, eBioscience, #14-5643), mouse anti-MyHC (1:2 MF20, DSHB), anti-mouse secondary antibody Alexa Fluor 555 conjugated (1:100, Life technologies A-21425) and anti-mouse secondary antibody Alexa Fluor 488 conjugated (1:100, Life technologies A-11001). Following the incubation with the secondary antibody, cells were washed two times with PBS and adipocytes were incubated with Oil Red O (Sigma #O0625) for 5 minutes. The samples were washed three times and nuclei were counterstained with Hoechst 33342 (Thermo Fisher Scientific, #3570) (1 mg/ml, 5 minutes at RT). Images were acquired with a LEICA fluorescent microscope (DMI6000B).

The total corrected cellular fluorescence (TCCF) was evaluated using ImageJ software (National Institutes of Health) as TCCF = ID – (ASC x MFBR). Where ID is integrated density, ASC is the area of selected cell and MFBR is the mean background fluorescence ^14^.

Microscope images have been processed changing only brightness and contrast and changes have been applied equally across the entire image and equally to controls.

### Cell differentiation

The percentage of myogenin positive cells (*Myog*^+^) was calculated as the ratio between the myogenin expressing nuclei and the total number of nuclei in each field.

The fusion index (*F_ind_*) was determined as the percentage of nuclei included in MyHC-expressing myotubes (containing at least 3 nuclei) *vs* the total number of nuclei ^15^.

The percent variance for *F_ind_* and for *Myog*^+^ are defined as

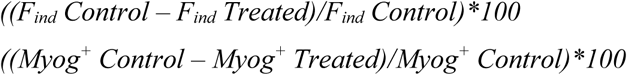

Myotube diameter was evaluated by taking three short-axis measurements at ¼, ½ and ¾ along the length of a given myotube and averaging them. More than 30 myotubes per condition were measured and data replicated in at least three independent experiments.

### RNA isolation, RNAseq and quantitative PCR

For qRT-PCR, total RNA was isolated from cells using Qiagen RNA Isolation Kit (#74106), RNA concentration, purity and integrity were measured in a spectrophotometer (NANODROP lite, Thermo SCIENTIFIC, Waltham, MA, USA). 0,5-1 μg were retrotranscribed using High-Capacity cDNA Reverse Transcription Kit (Applied Biosystems, # 4368814). Real time quantitative PCR was performed to analyze relative gene expression levels using SYBR Green Master mix 2X (Genespin # 44-QSTS-RSMMIX 200). Relative expression values were normalized to the housekeeping gene Tubulin.

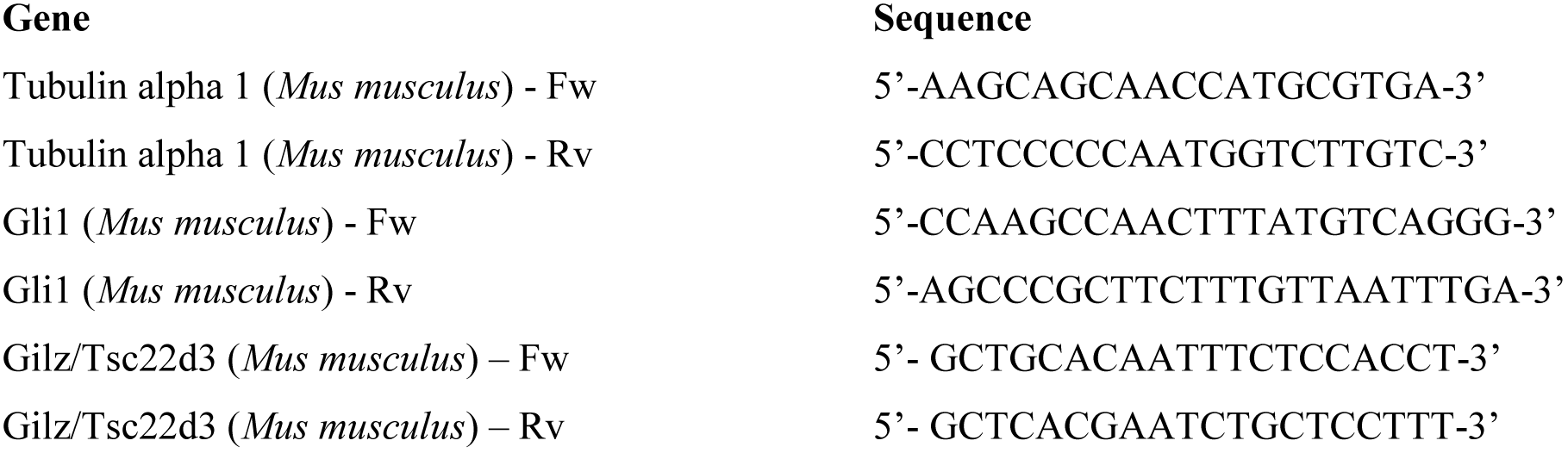

For RNAseq experiment, total RNA was extracted using Trizol from 3 independent preparations of control FAPs cells and 3 of cells treated with 5 μM budesonide for 24 hours in fGM. Libraries were prepared from 100 ng of total RNA using the QuantSeq 3’ mRNA-Seq Library Prep Kit FWD for Illumina (Lexogen GmbH). The library quality was assessed by using screen tape High sensitivity DNA D1000 (Agilent Technologies). Libraries were sequenced on a NextSeq 500 using a high-output single-end, 75 cycles, v2 Kit (Illumina Inc.). Approximately 44*10^6^ reads were obtained for each sample. Sequence reads were trimmed using the Trim Galore software ^16^ to remove adapter sequences and low-quality end bases (Q < 20). Alignment was performed with STAR ^17^ on the reference provided by UCSC Genome Browser ^18^ for *mus musculus* (UCSC Genome Build mm10). The expression levels of genes were determined with htseq-count ^19^ using the Gencode/Ensembl gene model. Differential expression analysis was performed using edgeR ^20^. Genes with a log2 expression ratio >|0.42| (treated/control sample) difference with a p-value < 0.05 and a FDR of < 0.1 were labeled as differentially expressed.

### Statistical analysis

Statistical analysis was performed by unpaired Student’s t test or One-way ANOVA (*p≤0.05, **p≤0,01, ***p≤0,001, ****p≤0,0001). Results are presented as the means ± SEM. All the experiments were repeated at least twice. p-values ≤ 0.05 were considered significant.

## Results

### Fluorescence microscopy-based screening of inhibitors of fibro-adipogenic differentiation

We developed a fluorescent microscopy-based protocol for the screening of compounds that modulate adipogenic differentiation of FAPs isolated from *mdx* mice, a model of Duchenne muscular dystrophy. FAPs were isolated, by magnetic bead separation, from 45-day-old *mdx* mice as CD31−/CD45−/ITGA7−/SCA1+ cells. Purified FAPs were plated in 384 well plates at a density of 1500 cells/well in GM containing 1 μg/ml of insulin. One day after plating each of the 1120 compounds of the Prestwick library were added at 5 μM final concentration and incubated for 6 additional days. Adipogenic differentiation was assessed by staining with ORO ^5^ (Fig. 1A). Compound cytotoxicity was assessed by counting Hoechst stained nuclei. DMSO 0.05% and TSA (20 nM) were used as negative and positive controls respectively. Among the compounds that we identified as active on adipogenesis, we noticed an enrichment of glucorticoids (GCs). GCs or structurally related steroid compounds represent the 7,5% of the screened drugs, while they are 24% in the antiadipogenic hit list. This corresponds to an enrichment factor of more than 3 (p = 0.02) and suggests a significant negative impact of glucocorticoids on the modulation of FAPs differentiation. The enrichment of glucocorticoids among the drugs that negatively affect adipogenesis came as a surprise, as glucocorticoids have been described as promoter of adipogenesis. This observation prompted us to investigate the underlying molecular mechanisms. For further characterization, we selected budesonide, clobetasol and halcinonide as being the GCs showing a high, intermediate and a low antiadipogenic activity in our assay (Table S1).

**Fig. 1.**
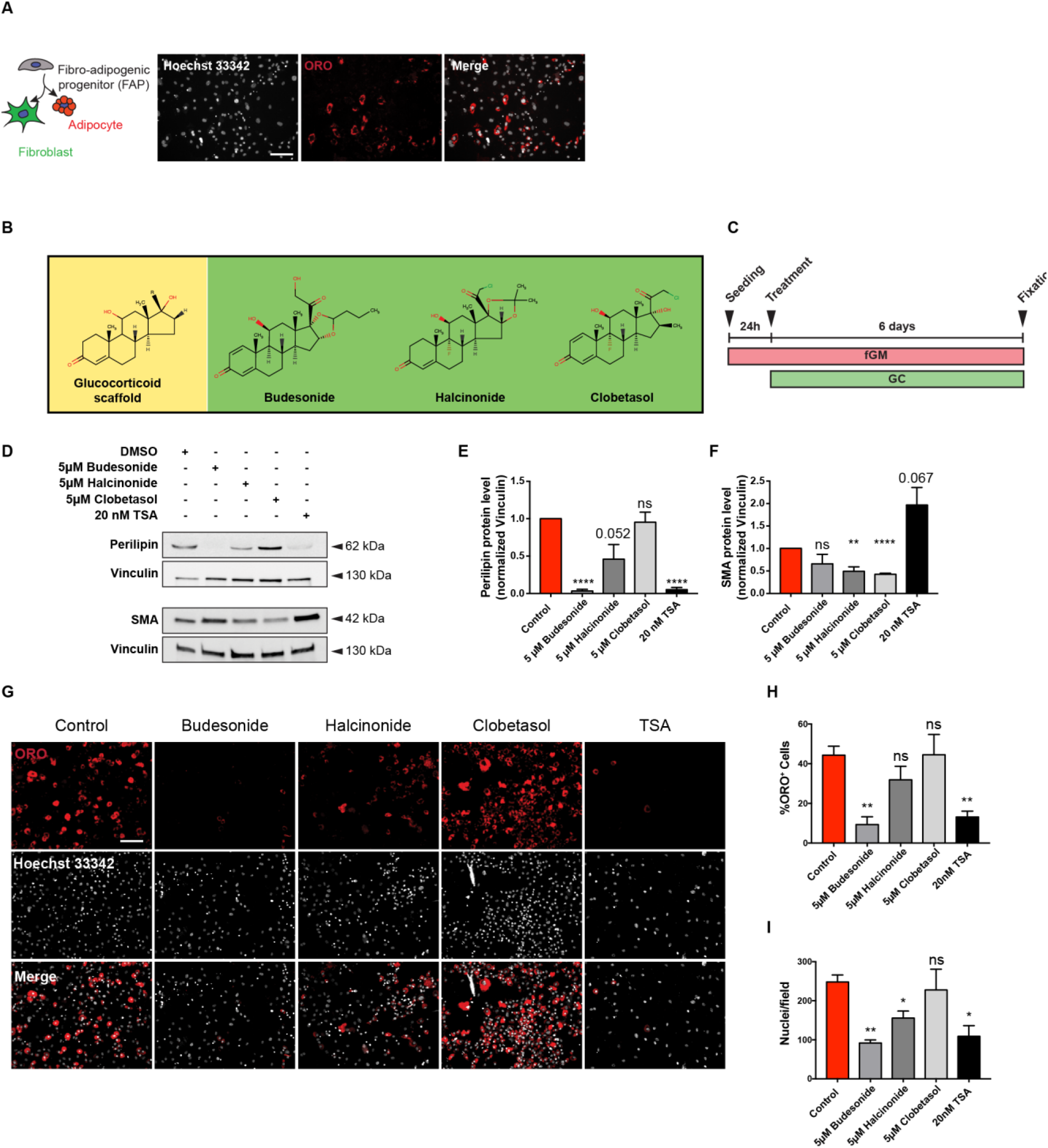
Budesonide affects FAPs adipogenic differentiation. (A) FAPs from *mdx* mice are incubated for 7 days in fGM. Cells were stained with ORO (red) to reveal adipocytes while Hoechst 33342 was used to stain nuclei (grey). (B) Structures of the GCs scaffold, budesonide and halcinonide and clobetasol. (C) Schematic representation of the experimental procedure for *mdx* FAPs treatment. FAPs were plated in fGM and after 24 hours cells were treated with 5 μM budesonide or halcinonide or clobetasol while TSA was used as positive control of adipogenic inhibition (D) Representative western blot showing perilipin and SMA expression in crude protein extracts from *mdx* FAPs cultured as reported in C. 30 μg of cell extracts were loaded in each lane. Vinculin is used as a loading control. (E, F) The bar graphs illustrate the densitometric quantitation of perilipin and SMA expression for the experiment reported in D. (G) Immunofluorescence images showing ORO (red) and Hoechst 33342 (gray) staining for FAPs treated as in C. (H, I) Bar plots showing the percentage of ORO positive cells and the number of nuclei/field for the experiment reported in G. The values are means of at least three independent experiments ± SEM. Statistical significance was evaluated using the Student’s t-test (*p≤0.05, **p≤0.01, ****p≤0.0001, ns: not significant). Scale bar: 100 μm.

Since FAPs are bipotent stem cells able to differentiate into both adipocytes and fibroblasts, we first aimed at confirming the impact of budesonide, halcinonide and clobetasol on adipogenic or fibrogenic differentiation of *mdx* FAPs. Perilipin ^21, 22^ and SMA expression were monitored by western blots as markers of adipogenesis and fibroblast differentiation respectively. Budesonide and TSA, and to a lower extent halcinonide, negatively affected perilipin expression at these concentrations (Fig. 1D, E) while treatment with halcinonide and clobetasol negatively affected expression of SMA (Fig. 1D, F). We further confirmed the effect of these compounds on *mdx* FAPs adipogenic differentiation by ORO staining (Fig. 1G, H).

GCs share the same cytosolic receptor and their different effects may be explained by a differential interaction with alternative distinct cellular targets. In this respect, it has been shown that some GCs also affect Smoothened (Smo) localization thereby modulating the activation of the sonic hedgehog pathway and inducing different phenotypes ^23, 24^. We observed increased levels of Gli1 mRNA, a downstream effector of the sonic hedgehog pathway, in 3T3-L1 cells treated with halcinonide, clobetasol and the Smo agonist SAG, while Gli1 expression did not change following treatment with budesonide or the Smo antagonist itraconazole (Fig. S1). Thus, despite being members of the same chemical class and activating the same receptor, budesonide halcinonide and clobetasol affect FAPs differentiation differently, possibly as a consequence of different modulation of alternative differentiation pathways.

### Budesonide affects PPARγ expression

Peroxisome proliferator-activated receptor γ (PPARγ) is the master regulator of adipogenesis. PPARγ expression is both necessary and sufficient for adipogenic differentiation ^25–27^. Freshly isolated FAPs do not express PPARγ and its expression increases during differentiation ^3, 4^.

We investigated whether budesonide impairs adipogenic commitment or rather compromises adipocyte maturation. To answer this question, we cultured *mdx* FAPs in fGM. 24 hours after seeding, cells were treated for further 6 days with 5 μM budesonide and PPARγ expression was assessed. As shown in Fig. 2A and B, *mdx* FAPs treated with budesonide, have a significant reduction in PPARγ expression suggesting an impairment of adipogenic commitment.

**Fig. 2.**
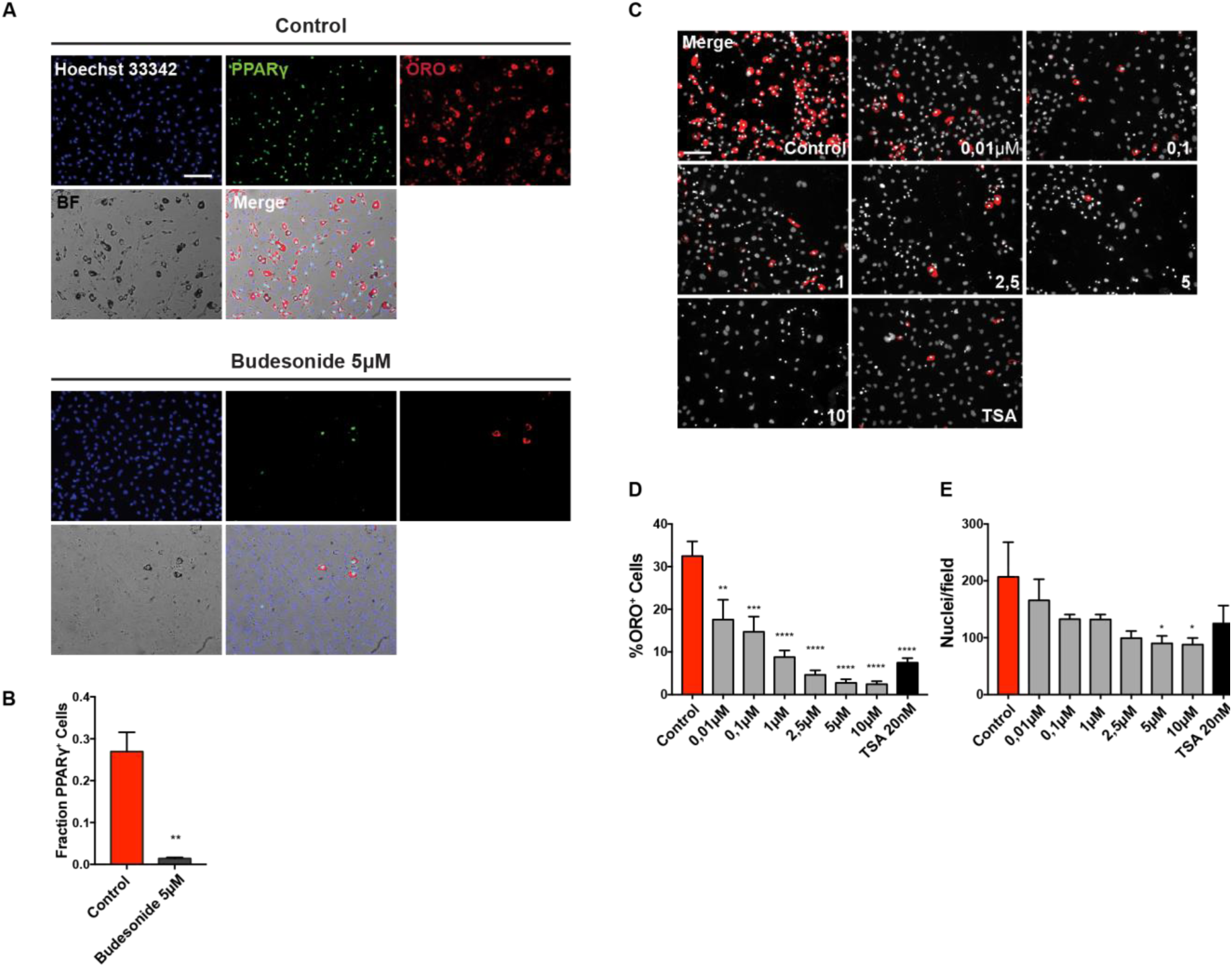
Budesonide inhibits PPARγ expression during *mdx* FAPs differentiation. (A) Immunofluorescence images showing *mdx* FAPs treated with 5 μM budesonide for 6 days and then stained with ORO (red) and an antibody against PPARγ (green). Nuclei are stained using Hoechst 33342 (blue) (B) Bar plot representing the fraction of PPARγ positive cells for the experiment reported in A. The values are means of three independent experiments ± SEM. Statistical significance has been evaluated using the unpaired T-Test (**p≤0.01). (C) Representative images of *mdx* FAPs treated with increasing concentrations of budesonide or with 20nM TSA. FAPs are stained with ORO (red) and Hoechst 33342 for nuclei (grey). (D) Bar plot showing the percentage of ORO positive cells for each concentration of budesonide. (E) The graph shows the number of nuclei in each field for controls and after treatment with budesonide. The values are means of three independent experiments ± SEM. Statistical significance has been evaluated using one-way ANOVA (*p≤0.05, **p≤0.01, ***p ≤0.001, ****p ≤ 0.0001, ns: not significant). Scale bar: 100 μm.

We next determined the dose response curve of budesonide treatment to evaluate its effective concentration for inhibition of adipogenic differentiation and toxicity. *Mdx* FAPs were isolated and allowed to differentiate with progressively higher concentrations of budesonide (ranging from 10 nM to 10 μM). As shown in Fig. 2C-F, budesonide significantly reduces the fraction of FAPs that differentiate into adipocytes already at 10 nM (Fig. 2D). The dose dependent negative modulation of adipogenesis is accompanied by a reduction in the number of nuclei at the end of the treatment.

### Budesonide stimulates terminal differentiation of satellite cells

To have a comprehensive view of a foreseeable impact of systemic budesonide treatment on muscle homeostasis we tested whether the drug has any effect on satellite cells, which play a prominent role in muscle regeneration ^28, 29^.

Satellite cells were purified from muscles of *mdx* mice as CD45−/CD31−/ITGA7+ cells by the magnetic bead technology. At day 5 post-treatment, the percentage of myogenin positive cells was significantly higher in samples treated with 1 and 5 μM of budesonide when compared to controls (Fig. 3A-C). In addition, we also observed an increase of MyHC (Myosin Heavy Chain) expression correlating with an increased fusion index and myotube diameter (Fig. 3D-F). We also observed increased myogenic differentiation upon treatment with budesonide in satellite cells isolated from wild type mice (Fig. S2)

**Fig. 3.**
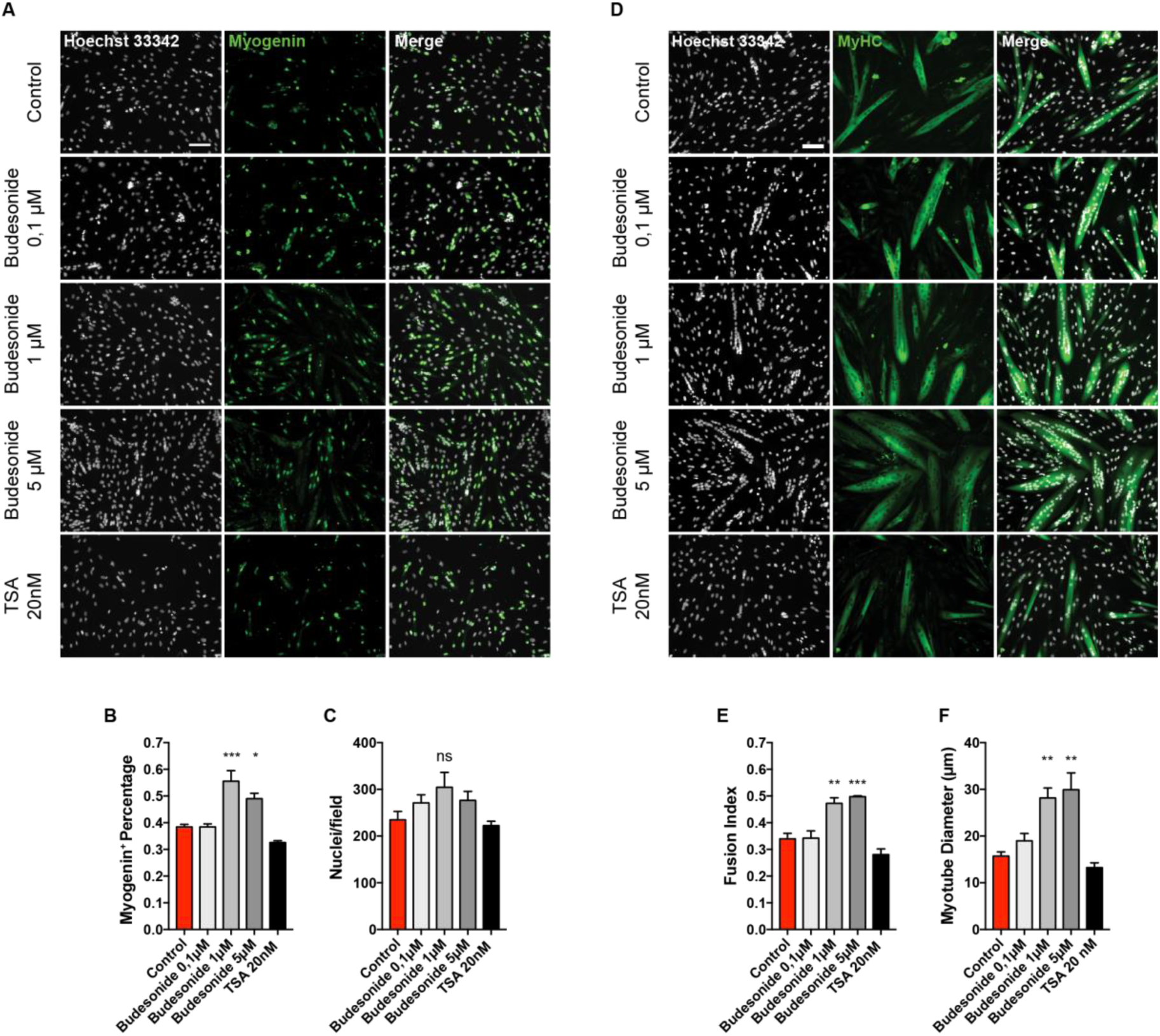
Budesonide treatment promotes terminal differentiation of *mdx* satellite cells. SCs were isolated from muscles of *mdx* mice as CD45−/CD31−/ITGA7+ cells and plated in sGM. 48 hours after plating, cells were treated with three concentrations of budesonide (0.1, 1 and 5 μM) or TSA (20 nm) for 5 additional days. Myogenic differentiation was assessed by immunostaining with antibodies against myogenin (A) and MyHC (D), early and late muscle-specific differentiation marker respectively. Nuclei were counterstained with Hoechst 33342. (B) Column chart showing the percentages of myogenin positive cells in the experiment in panel A. (C) Bar plot reporting the number of nuclei per field for the experiments in A and D. (E, F) Bar plots showing the fusion index and myotube diameter for the experiment in D. The values are mean of at least three independent experiments ± SEM. Statistical significance was evaluated using one-way ANOVA (*p≤0.05, **p≤0.01, ***P ≤0.001, ns: not significant). Scale bar: 100 μm.

### Inhibition of adipogenesis and stimulation of myogenesis by budesonide are both mediated by the glucocorticoid receptor

Most GCs effects are mediated by the glucocorticoid receptor (GCr) ^23, 30, 31^. We therefore asked whether the anti-adipogenic effect of budesonide is also mediated by the interaction with the GCr. As shown in Fig. 4A, B the incubation with the glucocorticoid antagonist mifepristone (RU-486) ^32, 33^ relieves the inhibitory effect of budesonide on FAPs differentiation (Fig. S3). We used dexamethasone as control for the activation of the GCr. The fraction of ORO positive cells following budesonide treatment was significantly reduced when compared to control and a similar effect was observed on cells incubated with dexamethasone. However, cells treated with RU-486 were largely insensitive to GCs-mediated inhibition of adipogenesis at 0,1 or 1 μM (Fig. 4A) suggesting that the anti-adipogenic effect of the two GCs is mediated by the GCr. We next asked whether the positive modulation of myogenesis is also mediated by the interaction of budesonide with the glucocorticoid receptor. To test this, 24 hours after seeding, C2C12 myoblasts were treated for 6 additional days with budesonide or dexamethasone. As observed in *mdx* satellite cells, treatment of C2C12 myoblasts with budesonide or dexamethasone induced an increase of the fusion index, paralleled by an increase of nuclei number (Fig. S4). In addition, as already observed for the anti-adipogenic effect, also the pro-myogenic effect is suppressed when glucocorticoids are administered in combination with the inhibitor of the glucocorticoid receptor RU-486 (Fig. 5A).

**Fig. 4.**
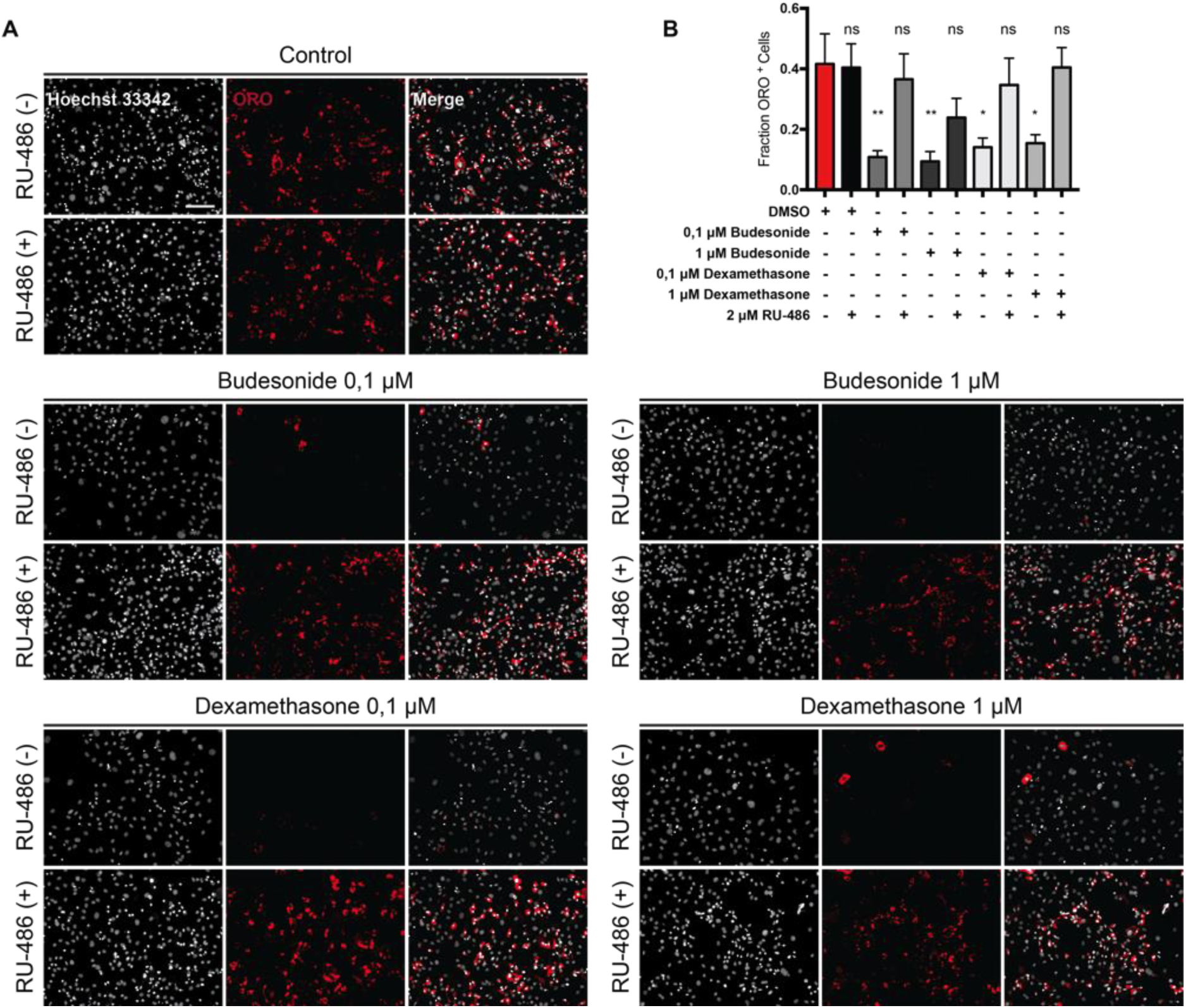
RU-486 counteracts budesonide or dexamethasone inhibition of FAPs adipogenic differentiation. *mdx* FAPs were isolated by the standard procedure and plated in fGM. After 24 hours, cells were treated with 0,1, 1 or 5 μM of budesonide or dexamethasone either with or without RU-486. After 6 days, cells were stained with ORO to evaluate adipocyte formation. (A) Immunofluorescence showing ORO staining (red) following *mdx* FAPs differentiation. Nuclei are stained with Hoechst 33342 and are shown in grey. (B) Bar plot showing the fraction of ORO positive cells. n=3-4 ± SEM. Statistical significance has been evaluated using one-way ANOVA (*p≤0.05, **p≤0.01, ns: not significant). Scale bar: 100 μm.

**Fig. 5.**
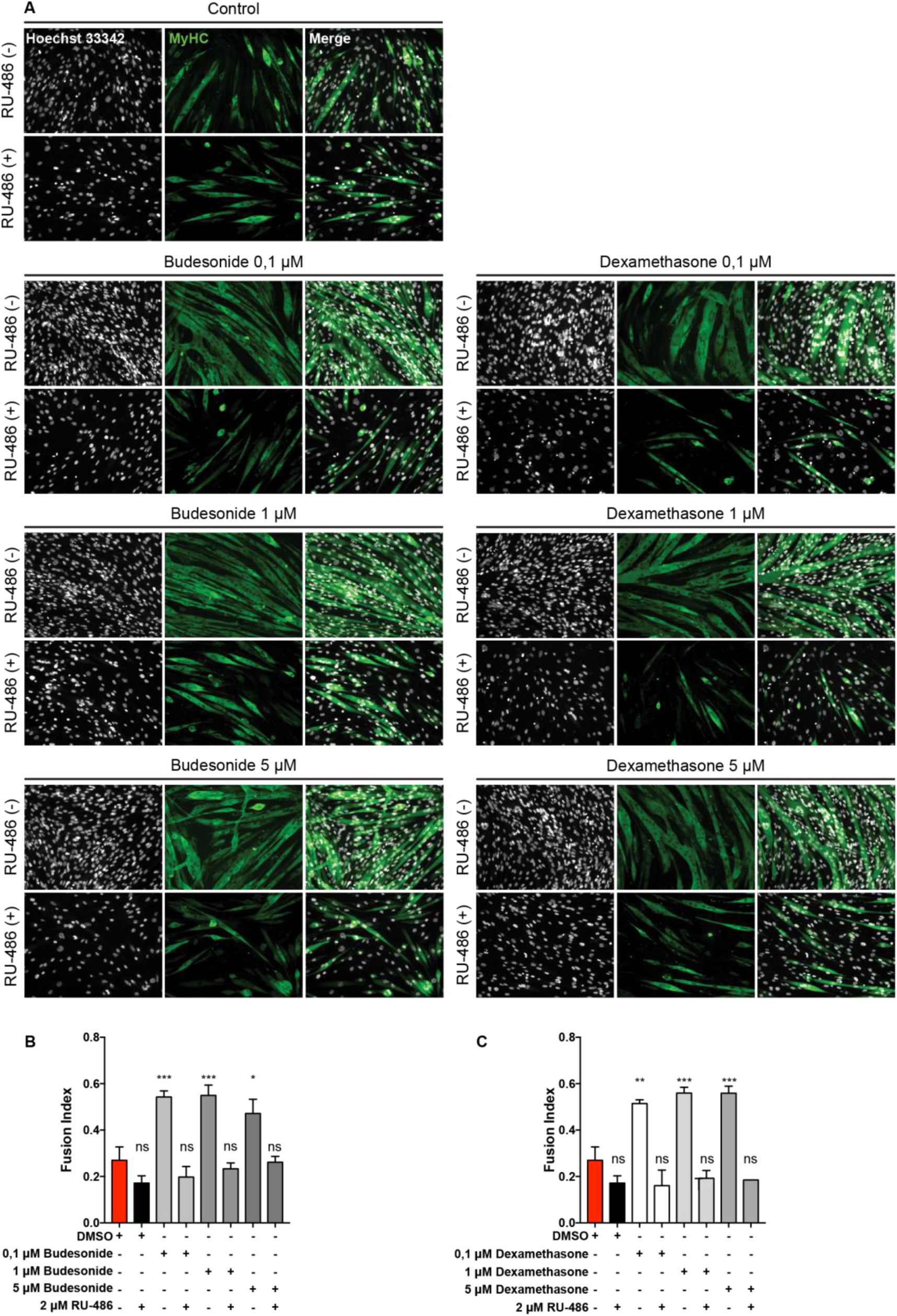
RU-486 suppresses the glucocorticoid-induced myogenic differentiation of C2C12 myoblast. (A) Immunofluorescence microphotographs of C2C12 labelled with an antibody against MyHC (green) after 7 days of culture in cGM. 24 hours after seeding, cells were treated with budesonide or dexamethasone at different concentrations alone or in combination with RU-486 (2μM) for 6 additional days. Nuclei are stained with Hoechst 33342 (grey). (B, C) Bar plots showing the fusion index for the experiment reported in A. n=4 ± SEM, Statistical significance has been evaluated using one-way ANOVA (*p≤0.05, **p ≤0.01, ***p≤0.001, ****p≤ 0.0001, ns: not significant). Scale bar: 100 μm.

### Budesonide can either act as a pro or anti-adipogenic drug depending on administration conditions

Dexamethasone is a glucocorticoid that promotes terminal differentiation of pre-adipocyte ^34, 35^. This is in contrast with the anti-adipogenic effect that we observe for budesonide and dexamethasone when administered to FAPs in our screening conditions. Standard differentiation protocols for pre-adipocytes such as 3T3-L1 include the expansion of pre-adipocytes *in vitro* and their incubation for 48 hours after reaching confluence and before switching to adipogenic induction medium (AIM) containing 1 μg/ml insulin, 0.5 mM IBMX and 1 μM dexamethasone. After 48 hours in AIM, cells are exposed to adipogenic maintenance medium (MM) containing 1 μg/ml of insulin ^36^ for two additional days. We therefore wondered if the anti-adipogenic effect of budesonide as observed in sub confluent FAPs was also present if FAPs were induced to differentiate according to the “standard” protocol. To address this point, freshly isolated *mdx* FAPs were cultured in fGM in the absence of insulin for 6 days. Confluent cells were next treated with budesonide or dexamethasone alone or in combination with the AIM pro-adipogenic components, insulin and IBMX, for two days. After 48 hours, cells were switched to MM for 48 additional hours. When FAPs are treated according to this protocol and reach confluence in the absence of adipogenic stimuli they differentiate poorly. In these conditions the inhibitory effects of budesonide or dexamethasone on this low basal differentiation level are difficult to measure. Conversely, if switched to AIM, confluent FAPs differentiate more efficiently and the addition of glucocorticoids to the adipogenic mix, differently from what was observed on freshly isolated FAPs, increases adipogenic differentiation. We wanted to exclude that the observed anti-adipogenic effect on freshly isolate FAPs was an artefactual consequence of the stress caused by the purification procedure. To address this point, we first allowed freshly purified FAPs (P0) to recover for four days in a commercial growth factor-rich medium (Cytogrow). Cells were then collected and plated (P1) in fGM at the density of (300 cells/mm^2^). Budesonide was added after 24 hours and cells were incubated for 6 additional days. Similarly to FAPs P0, also FAPs P1 maintain sensitivity to the anti-adipogenic effect of budesonide (Fig. 6D, E). We conclude that budesonide exerts a significant anti-adipogenic activity when FAPs are treated while they are actively growing and before they reach confluence and become insensitive to budesonide inhibition. Similarly, to what has been reported in the literature for other glucocorticoids ^37^ budesonide, at cell confluence, promotes adipogenesis only if administered in addition to the standard components of the adipogenic mix.

**Fig. 6.**
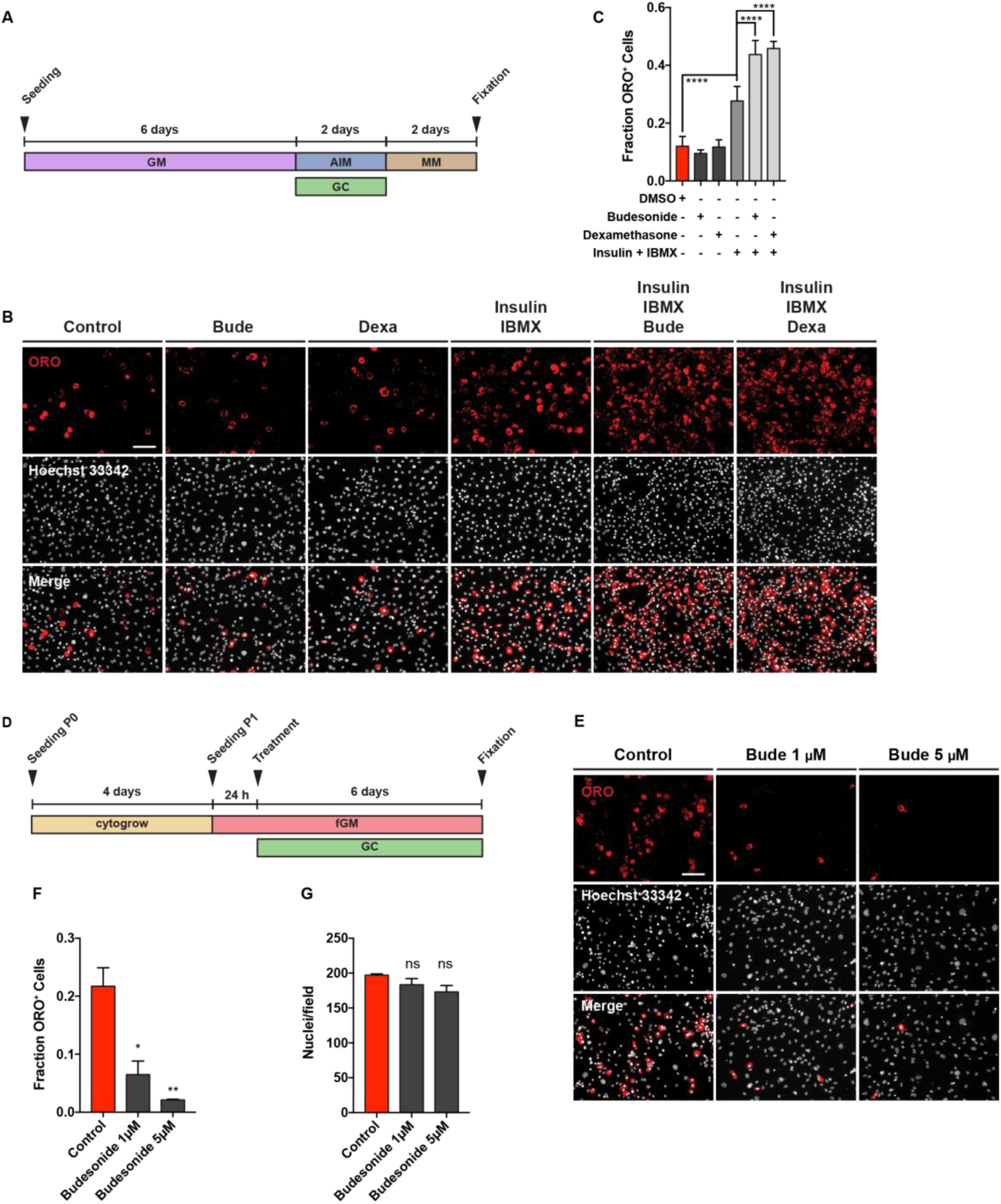
Different effect of GCs on adipogenic differentiation depending on the timing of their administration. (A) Schematic representation of the experiment reported in B, *mdx* FAPs were isolated by the standard procedure and plated in GM. 6 days post-seeding, confluent cells were treated with 1 μM of budesonide or dexamethasone either with or without the AIM (10% FBS, 1 μg/ml of insulin, 0.5 mM IBMX) for 2 days. Cells were then moved to MM (10% FBS and 1 μg/ml of insulin) and incubated for 2 additional days. (B) Immunofluorescence microphotograph of cells stained with ORO to reveal adipocyte formation (red) and Hoechst 33342 (grey). (C) Bar plot showing the fraction of ORO positive cells for the experiment reported in panel B. n=3-4 ± SEM. Scale bar: 100 μm. (D) Schematic representation of the experiment reported in E, FAPs isolated from young *mdx* mice were expanded in Cytogrow for 4 days until they reached 70% confluence. Cells were then detached and plated in fGM. 24 hours after seeding cells were treated with 1 μM or 5μM budesonide for 6 days. (E) FAPs adipogenic differentiation was assessed by staining cells with ORO (red) and Hoechst 33342 (grey). (F, G) Bar graphs presenting the quantitation of the adipogenic differentiation and the number of nuclei per field for the experiment in panel E. n=2 ± SEM. Statistical significance tested by one-way ANOVA (*p≤0.05, **p ≤0.01, ****p≤ 0.0001, ns: not significant). Scale bar: 100 μm.

### cAMP modulation affects the anti-adipogenic effect of budesonide on sub-confluent FAPs

Since we observed that GCs have a pro-adipogenic activity when confluent *mdx* FAPs are exposed to specific GCs in combination with the adipogenic induction medium, we wondered if this was true also on sub-confluent FAPs. To answer this question, we plated FAPs in GM alone or supplemented with insulin, IBMX or both. 24 hours after plating, cells were treated with increasing concentrations of budesonide. In these culture conditions the adipogenic differentiation of cells incubated in GM alone or supplemented with insulin are sensitive to the anti-adipogenic effect of budesonide (Fig. 7A-D). By contrast, cells incubated in media supplemented with IBMX, either alone or in combination with insulin, are markedly less sensitive to inhibition of adipogenesis (Fig. 7E-H). Beside the effect on adipogenic inhibition, IBMX treatment is also associated with a significant decrease of nuclei number when compared to cell maintained in GM or GM supplemented with insulin (Fig. S5A, B). Since IBMX is a non-competitive inhibitor of phosphodiesterase we hypothesized that an increase of the intracellular levels of cAMP could be the cause of the insensitivity to budesonide inhibition. To test this hypothesis, we incubated FAPs with forskolin, an activator of adenylyl cyclase also causing an increase in the levels of cAMP. Similarly, to what observed in IBMX treated cells, the fraction of ORO positive cells is not significantly reduced when cells are treated with budesonide in combination with forskolin (Fig. 7I, L). However, although forskolin was efficient in relieving the antiadipogenic effect of budesonide, as monitored by the fraction of ORO positive cells (Fig. 7L, M), differently from IBMX treatment we observed that, in these conditions, adipocytes are less mature and are characterized by a lower intensity of ORO staining (see insets of Fig. 7I, Fig. S5C, D). We conclude that an increase of cytosolic cAMP is epistatic on the budesonide capacity to negatively affect adipogenesis, independently of the proliferative condition of the cell.

**Fig. 7.**
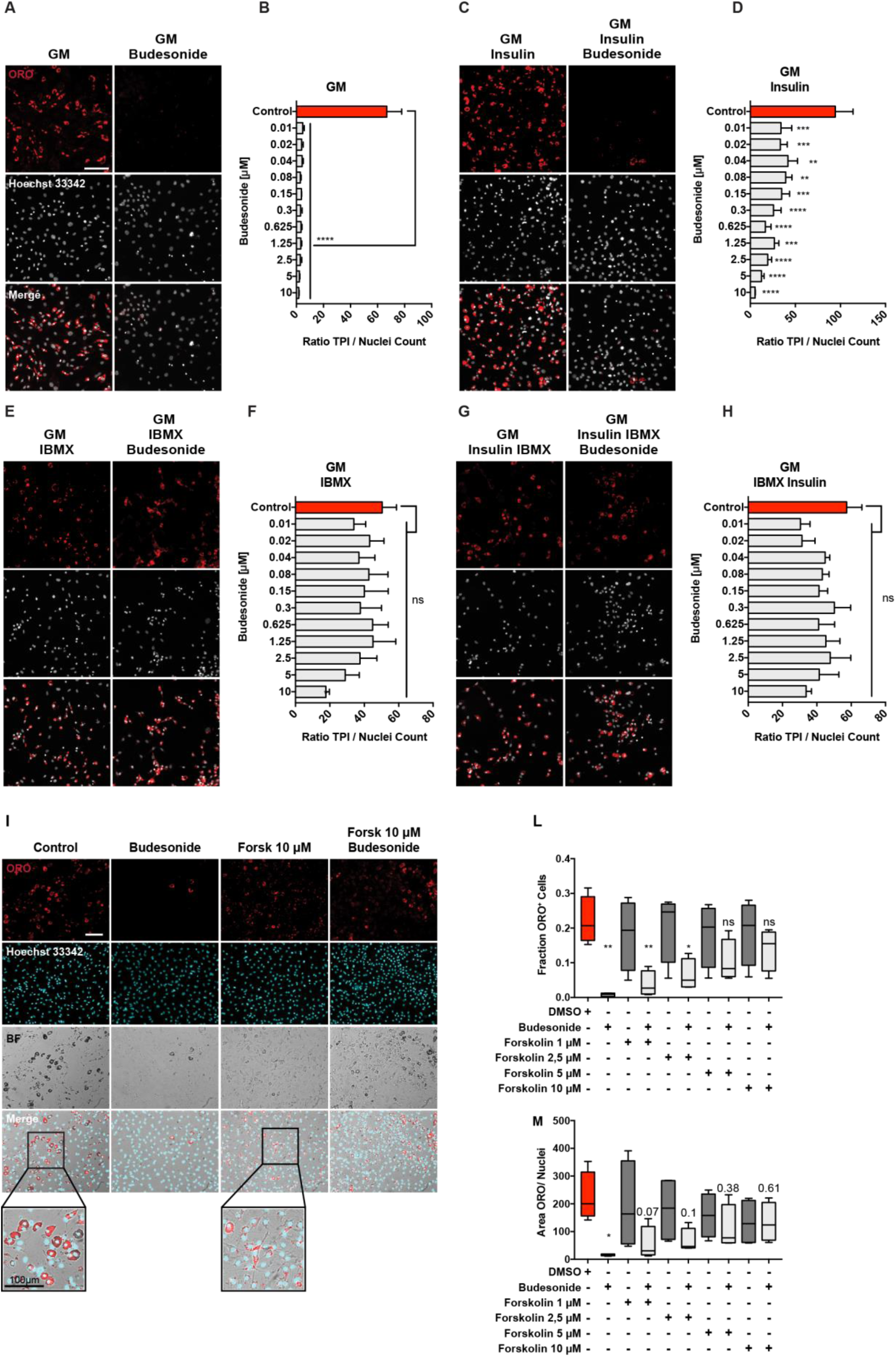
Increasing cAMP levels contrasts the anti-adipogenic effect of budesonide. The panel shows *mdx* FAPs isolated by the standard procedure and plated in GM (A) or supplemented with 1 μg/ml insulin (C), 0.5 mM IBMX (E) or insulin and IBMX (G). 24 hours after plating cells were treated with increasing concentration of budesonide for further 6 days. Cells were stained with ORO (red) to identify adipocytes while Hoechst 33342 was used for nuclei counterstain. The bar plots indicate the ratio between the total pixel intensity (TPI) and the total number of nuclei for the different concentrations of budesonide in each culture condition: GM (B), GM + insulin (D), GM + IBMX (F) and GM + insulin + IBMX (H). Values are the means of three different experiments ± SEM. (I) Immunofluorescence images showing *mdx* FAPs isolated by the standard procedure and plated in fGM. 24 hours upon seeding, cells were treated with increasing concentrations of forskolin in presence or absence of budesonide 5 μM. Adipogenic differentiation was assessed using ORO staining to reveal adipocytes and Hoechst 33342 to reveal nuclei. The insets display a higher magnification of the merged channels. (L, M) box plot showing the fraction of ORO positive cells or the ratio between the area covered by ORO positive signal and nuclei for the experiment reported in I. Box plots show median and interquartile range with whiskers extended to minimum and maximum values. In M the reported values represent the p-value. n=3. Statistical significance has been evaluated using one-way ANOVA (*p≤0.05, **p ≤0.01, ***p≤0.001, ****p≤ 0.0001, ns: not significant). Scale bar: 100 μm.

### Pro or anti-adipogenic effects of budesonide correlate with Gilz expression

To gain insights into the mechanisms underlying the observed inhibition of adipogenesis by budesonide we performed an RNAseq experiment to identify genes whose expression is perturbed by drug treatment (Fig. S6). We identified transcripts for a total of 14381 genes: 87 genes were significantly up-regulated while 79 were down-regulated by budesonide treatment (Table S2). By entering these lists of modulated genes in the DAVID online tool ^38^ did not reveal any significant enrichment in gene ontology annotation or KEGG pathways after correction for multiple testing. However, by inspecting the list of genes that were significantly upregulated we noticed that the fifth most upregulated gene was Gilz (Tsc22d3) (14x fold change), which encodes an established antagonist of the PPARγ transcription factor.

The glucocorticoid-induced leucine zipper (Gilz/TSC22D3) is a primary target of glucocorticoids/GCr and a known mediator of the anti-inflammatory, immunosuppressive, and anti-proliferative actions of glucocorticoids in many cell types ^39, 40^. Gilz antagonizes adipocyte differentiation of mesenchymal stem cells by binding to the PPARγ2 promoter and inhibiting its transcription ^41^. To confirm that Gilz was involved in adipogenesis inhibition of sub-confluent FAPs mediated by budesonide we monitored Gilz mRNA and protein levels at 24 and 48 hours after budesonide treatment (Fig. 8A). After 24 hours, Gilz mRNA levels are significantly upregulated (approximately 30 folds) compared to control. This is paralleled by an increase in the protein level at both time points (Fig. 8C, E, G). No equivalent upregulation of Gilz mRNA or protein levels were observed when cells were treated with glucocorticoids according to the “standard” *ex vivo* adipogenesis induction protocol (i.e., IBMX and Insulin) (Fig. 8B, D, F, H). These observations point to Gilz expression, mediated by the activation of the GCr, as an essential step in the inhibition of adipogenesis mediated by glucocorticoids. We also analyzed our gene expression data by the web tool eXpression2Kinases X2K ^42^ that computes enrichment for modulated genes that are enriched in transcription factors binding sites. Interestingly 3 of the 6 transcription factors, or chromatin modifiers, with lowest p-value have already been implicated in the modulation of adipogenic differentiation (CEBPd, SUZ12, EP300, SOX9).

**Fig. 8.**
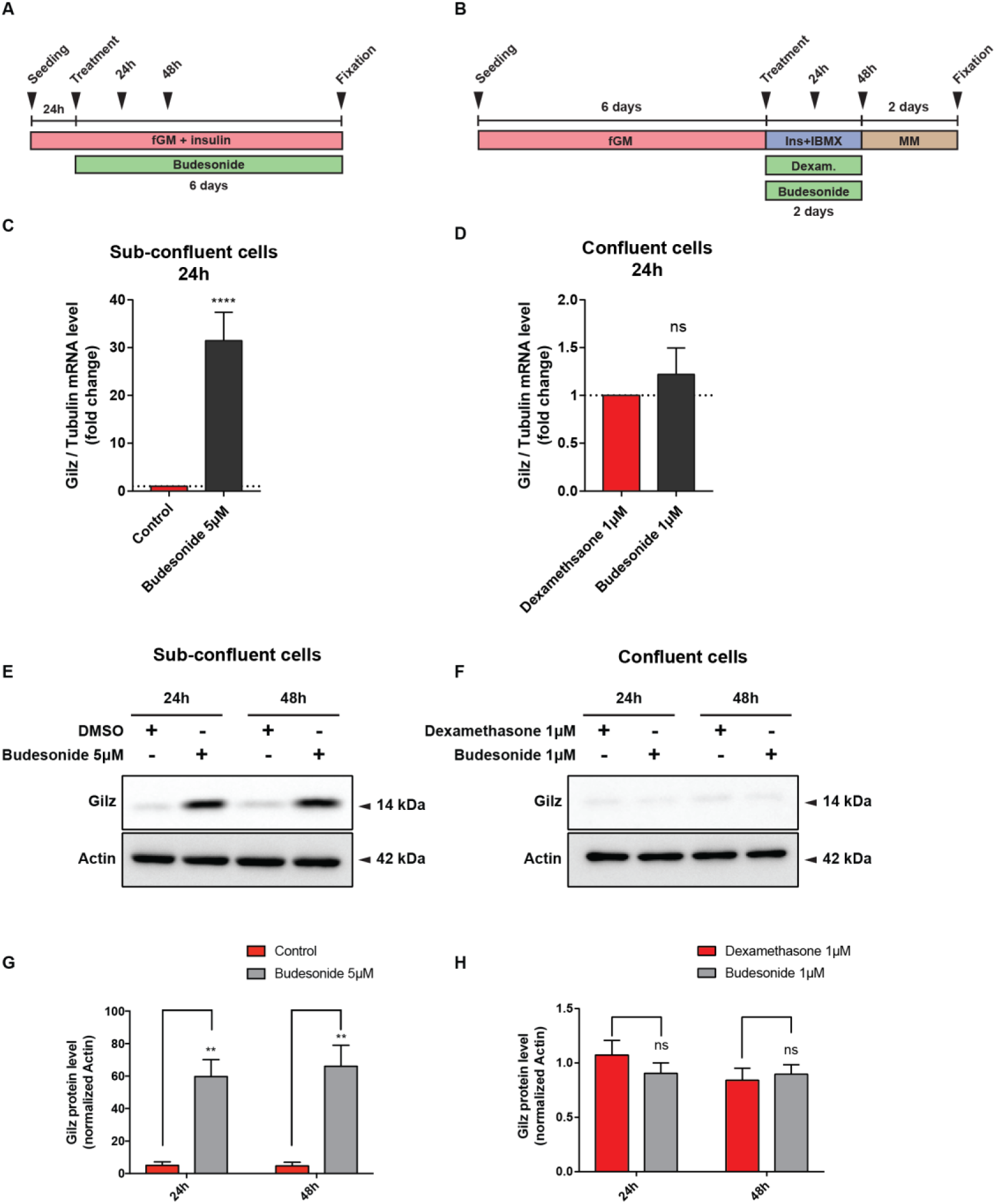
Budesonide induces Gilz expression only in sub-confluent FAPs. (A) Schematic representation of the experiments reported in C, E, G. *Mdx* FAPs isolated by the standard procedure were plated in fGM. 24 hours after plating, sub-confluent cells were treated with 5 μM of budesonide for further 6 days. (B) Schematic representation of the experiments reported in D, F, H. Mdx FAPs isolated by the standard procedure were plated in GM and after 6 days, confluent cells were exposed to the AIM complemented with dexamethasone 1 μM or budesonide 1 μM for further 2 days. (C, D) Bar plots showing the mRNA level of Gilz analyzed by RT-qPCR following 24 or 48 hours of treatment with budesonide for the conditions described in A and B. Tubulin was set as reference gene. (E, F) Immunoblot analysis revealing the protein content of Gilz after 24 and 48 hours of treatment with budesonide for the experimental conditions described in A and B. Actin is used as loading control. (G, H) Bar plots representing the quantitation of Gilz protein levels for the experiment reported in E and F. n=3 ± SEM. Statistical significance has been evaluated using two-way ANOVA (**p ≤0.01, ****p≤ 0.0001, ns: not significant).

## Discussion

In muscular dystrophy the degeneration of the muscle tissue is initially compensated by efficient regeneration counterbalancing muscle loss ^1^. However, over time, this process is impaired and myofiber repair is thwarted by the formation of fibrotic scars and fat infiltrations, undermining muscle function ^2^. Fat deposition and fibrosis are aggravating consequences of a failure in the mechanisms controlling the differentiation potential of fibro-adipogenic progenitors ^43^ Learning to control FAPs differentiation may help establishing therapeutic strategies to limit or delay excessive fat deposition and fibrosis associated with degenerative pathologies. To expand our pharmacological toolbox, we have devised a screening strategy aimed at identifying small molecules controlling adipogenic differentiation of FAPs purified from a dystrophic mouse model. Somewhat unexpected, the screening hit list was highly enriched in glucocorticoids, a class of molecules that have been shown to promote adipogenesis of mesenchymal stem cells ^37, 44^.

Interestingly, steroids represent the standard palliative treatment to slow down the progression of muscle degeneration and preserve muscle strength in DMD patients ^45–47^. Most of the clinical indications of GCs are related to their immune-modulating and anti-inflammatory effects ^48, 49^. However, glucocorticoids are highly pleiotropic molecules, affecting the physiology of practically any organ ^50^ and long term systemic administration is often accompanied by unwanted side effects ^48, 51–53^. The molecular and physiological mechanisms underlying their mild beneficial effects on dystrophic patients, and adverse side effects, are poorly understood ^54, 55^. Prompted by the results of our screening we have characterized the effects of glucocorticoid treatment on two primary muscle progenitor cell types, fibro-adipogenic progenitors and satellites cells.

Numerous, sometimes contradictory, reports implicate GCs in the modulation of differentiation of mesenchymal stem cells *in vivo* and in *vitro* (for a review see ^56^). However, the variability of the experimental conditions, including species, tissue source, plating density, passage number and culture conditions, hampers the definition of a clear picture. Focusing on adipogenesis, most reports demonstrate that GCs have a pro-adipogenic effect and weight gain is one of the most common side effects of prolonged GCs treatment ^57^. *In vitro* GCs promote adipogenic differentiation of mesenchymal stem cells ^58, 59^. However, it has also been reported that GCs may have an inhibitory effect on adipogenic differentiation ^60, 61^. In addition and relevant for their impact on treatment of muscle disorders glucocorticoids inhibit myogenesis by inducing the expression of the glucocorticoid-induced leucine zipper (Gilz) which in turn inhibits MyoD function ^62^. On the other hand, somewhat in contrast, the exposure of C2C12 myoblasts to dexamethasone causes an increase in proliferation rate and terminal myogenic differentiation ^63^. Finally, dexamethasone administration to myotubes produces an atrophic effect with increased expression of atrogin-1 and a decreased protein content of MyHC ^64–66^. Reconciling the conclusions of different reports is made difficult by the heterogeneity of GCs activity especially when tested on different cell types, primary or stable cell lines.

Although in our assay GCs, as a chemical class, showed a clear propensity to inhibit FAPs differentiation they also showed remarkable heterogeneity. Among the GCs that in the screening showed inhibitory activity on *mdx* FAPs differentiation, we further characterized budesonide, halcinonide and clobetasol.

Budesonide, halcinonide and clobetasol, despite sharing the same cytosolic receptor, modulate FAPs differentiation differently. Budesonide, as dexamethasone, significantly reduces adipogenic differentiation while FAPs exposure to halcinonide and clobetasol significantly decreases the expression of smooth muscle actin, a marker of fibrogenesis. This is possibly the consequence of differential interaction with additional cellular targets ^67^. For instance distinct GCs can differentially modulate the localization of the Shh target Smo ^23, 24, 27^. We observed that the mRNA levels of Gli1, a downstream effector of Shh is increased in preadipocyte 3T3-L1 treated with halcinonide and clobetasol and not with budesonide (Fig. S1). Overall these results suggest that the differences observed in the capacity of GCs to modulate fibro-adipogenic differentiation, may be related to their differential modulation of distinct cellular target besides the GCr. Our screening readout is based on the identification of lipid droplets, a rather late stage of adipocyte differentiation. However, the observation that budesonide negatively affects PPARγ the master regulator of adipogenic differentiation suggests that this glucocorticoid negatively modulates a relatively early step of the adipogenic commitment rather than a late differentiation step. Our conclusions are based on treatment of purified primary cells. Their *in vivo* relevance under standard therapeutic regimens can be estimated from the available pharmacological data. As shown here, budesonide activity on FAPs differentiation *in vitro* has a potency in the mid nanomolar, concentration range which is comparable with the range of plasma concentrations in patients treated with therapeutic glucocorticoid dosages ^68^.

The effect of GCs on fiber size homeostasis and myogenesis is also controversial. It was proposed that steroids could exert their beneficial effects on DMD patients by inhibiting muscle proteolysis ^69^. However, dexamethasone administration to myotubes produces an atrophic effect with increased expression of atrogin-1 ^64–66^ and muscle atrophy is one of the main side effects of prolonged GCs treatment ^70^.

In addition, several GCs such as dexamethasone and prednisolone, can exert positive or negative effects on myogenic cell lines, depending on the stage of administration. The exposure of C2C12 myoblast to dexamethasone causes an increase in terminal differentiation ^63^. Similarly, we observed that budesonide treatment of primary *mdx* satellite cells also promotes terminal differentiation. This could suggest that one potential beneficial effect of GCs treatment of DMD patients is the promotion of a more robust muscle regeneration. This beneficial effect, however, would be overridden by atrophy associated to the long-term exposure. It has been recently suggested that intermittent, rather than daily glucocorticoid administration could promote repair upon injury avoiding the side effect of atrophy induction ^49^. Our results support the notion that distinct GCs may exert different effects on muscle progenitor differentiation and suggest that in the choice of glucocorticoids to be used in DMD treatment secondary effects on muscle progenitor cells, other that their immunosuppressive properties, should be considered.

We have shown here that the “Janus-like” effect of GCs on mesenchymal stem cell adipogenic differentiation is not only a consequence of different response of different cell systems to GCs but can be reproduced when homogeneous primary cells are treated in different conditions. While glucocorticoid treatment of sub-confluent and actively growing FAPs cells has an important anti-adipogenic effect, treatment of confluent cells stimulates adipogenesis.

A second strong conclusion is that two positive modulators of cAMP levels, IBMX and forskolin, counteract the anti adipogenic effect of budesonide on sub-confluent FAPs. Our results suggest that budesonide and other glucocorticoids, including dexamethasone, can play a double edge game on muscle FAPs differentiation as they can promote or interfere with adipogenesis, depending on the different growing conditions.

We have shown that the effect of glucocorticoids on fibro-adipogenic progenitors correlates with the induction of transcription of the Gilz gene. Gilz is positively regulated by the GCr, plays a role in the anti-inflammatory and immunosuppressive effects of glucocorticoids ^71^ and is a potent inhibitor of adipogenesis induced by the GCr via inhibition of the PPARγ2 gene ^41^. It remains to be established why GCs administration does not lead to Gilz expression and adipogenesis inhibition in confluent preadipocytes or when the progenitor cells are treated with drugs that promote the accumulation of cAMP. The potency of GCr as a transcription factor is known to be modulated by several co-activators and co-repressors. Which of these are responsible for the reported differential pro- or anti-adipogenic effect in different experimental conditions requires further investigation.

We report here that distinct GCs can modulate the differentiation potential of two cell types critically involved in the regenerating muscle environment of a dystrophic mouse model. GCs can modulate both *mdx* FAPs adipogenic potential and *mdx* SC myogenic differentiation. Altogether, the results reported here suggest that GCs may exert their beneficial effect on DMD patients not only through the reduction of the inflammatory environment associated with the chronic DMD-associated muscle degeneration, but also through the modulation of stem cell differentiation. As we have shown that distinct GCs have different abilities to modulate FAPs differentiation and that their effect is dependent on whether the target cells are actively growing or have reached confluence and stopped cycling, it is difficult to predict how GCs that are currently used to treat muscular dystrophies are impinging on FAPs plasticity while these progenitor cells cycle between the proliferative and resting state that characterize DMD.

## Author contributions

The experiments presented in this manuscript have been conceived and planned by ACP, AR, GG, MC, AB, LC and GCs. AB designed the high content screening assay funnel and interpreted the screening data. MC run the screening designed the image analysis and contributed to compound identification. MS and SH designed the collection and the molecular features of the identified compounds. ACP, AR, GG, RS performed the experiments to validate GCs activity on all the cell type presented in the manuscript. ACP, AR, MC, GG, MS, RS, AP, SH, LC, AB and GCs contributed to the interpretation of the results. AP performed the RNAseq analysis. ACP, LC, and GCs wrote the manuscript with support from AR, GG and AB. LC, GCs and AB supervised the project.

## Acknowledgements

This work was supported by a grant of the European Research Council (grant N. 322749) to G.C. This work was made possible at IRBM by the CNCCS s.c.a.r.l. initiative

