## Supplementary material for "Janus effect of glucocorticoids on differentiation of muscle fibro/adipogenic progenitors"

Correspondence should be addressed to:

**Supplementary Figures Legends**

**Fig. S1. The expression of Gli1 is induced in 3T3-L1 treated with halcinonide and clobetasol.** 3T3-L1 pre-adipocytes were expanded *in vitro* and incubated as a confluent culture for 48h. Cells were then switched to AIM for further 48 h and treated with budesonide, halcinonide and clobetasol while unstimulated cells were kept in pre-adipocyte expansion medium. The bar plot represent Gli1 expression determined by quantitative PCR. n=3 ± SEM. SAG and Itraconazole were used as positive and negative control for Gli1 induction respectively. Tubulin was used as housekeeping gene. SEM is reported. Statistical significance tested by one-way ANOVA (*p≤0.05, **p ≤0.01, ****p≤ 0.0001, ns: not significant).

**Fig. S2. Budesonide treatment promotes terminal differentiation of wt satellite cells.** SCs were isolated from muscles of C57BL/6 mice as CD31-/CD45-/a7Integrin+ cells and plated in sGM (DMEM, 20% FBS, 10% horse serum, 1% Chicken Embryo Extract). 48 hours after plating, cells were treated with three concentrations of budesonide or dexamethasone (0.1, 1 and 5 μM) for 5 additional days. Myogenic differentiation was assessed by immunostaining with antibodies against Myosin Heavy Chain (MyHC) while nuclei were stained with Hoechst 33342. The bar plots show the fusion index (A), the percentage of the field area covered by myotubes (B) and the number of nuclei per field for the experiments (C). The values are mean of two independent experiments ± SEM. Statistical significance was evaluated using one-way ANOVA (*p≤0.05, **p≤0.01, ns: not significant).

**Fig. S3. RU-486 counteracts the reduction of nuclei number associated with budesonide treatment of *mdx* FAPs**. *mdx* FAPs were isolated by the standard procedure and plated in fGM. After 1 day, cells were treated with 0,1, 1 or 5 μM of budesonide or dexamethasone either with or without RU-486. After 6 days, cells were stained with ORO to evaluate adipocyte formation. (A) Bar plot showing the average number of nuclei per field. n=3-4 ± SEM. Statistical significance has been evaluated using one-way ANOVA (*p≤0.05, **p≤0.01, ns: not significant).

**Fig. S4 RU-486 counteracts the increase of nuclei number that follows the treatment of C2C12 myoblast with budesonide and dexamethasone.** C2C12 myoblast were plated in cGM. 24 h after seeding cells were treated with budesonide or dexamethasone at different concentrations alone or in combination with RU-486 (2μM) for 6 additional days and subsequently stained with an antibody against MyHC while nuclei were counterstained with Hoechst 33342. Bar plots showing the nuclei number per field for cells treated with budesonide (A) or dexamethasone (B). n=4. Statistical significance has been evaluated using one-way ANOVA (***p≤0.001, ****p≤ 0.0001, ns: not significant).

**Fig. S5. Treatment with IBMX or forskolin affect nuclei number or adipogenic differentiation of *mdx* FAPs.** *mdx* FAPs isolated by the standard procedure and plated in GM 20% FBS were supplemented with

0.5 mM IBMX 48 h after their seeding and cultured for further 5 days. (A) The microphotographs show nuclei stained with Hoechst 33342. (B) The bar plot indicates the average number of nuclei per field. Scale bar: 100 μm. (C) *mdx* FAPs were isolated by the standard procedure and plated in fGM. 24h upon seeding, cells were treated with increasing concentrations of forskolin in presence or absence of budesonide 5 μM for further 6 days. Adipogenic differentiation was assessed using oil red o staining to reveal adipocytes and Hoechst 33342 to reveal nuclei. The plot shows the percentage of adipocytes for a specific range of total corrected cellular fluorescence (TCCF) intensity value (expressed in arbitrary units AU) for control cells or cells treated with forskolin 10 μM alone or in combination with budesonide 5 μM. (D) Box plot showing the nuclei number per field. Box plots show median and interquartile range with whiskers extended to minimum and maximum values. n=3. Statistical significance has been evaluated using one-way ANOVA (*$p \leq 0.05$, **$p \leq 0.01$, ns: not significant).

**Fig. S6. Transcriptome analysis of *mdx* FAPs treated with budesonide.** (A) Multi-scatter plot of budesonide treated and control profiles, which highlights higher correlation between budesonide treated samples (mean correlation coefficient 0.9405) and control samples (mean correlation coefficient 0.963) in comparison with budesonide-treaded VS control samples (mean correlation coefficient 0.914). (B) The principal component analysis reveals a separation of budesonide-treated (blue area) and control samples (read area). (C) Density scatter plot of the expression in CPM (counts per million) of all genes of budesonide and control samples. Black dots represent genes whose enrichment id statistically significant genes. (D) Heatmap and hierarchical clustering of the significant (FDR < 0.05) differentially expressed genes in the two conditions. (E) Table showing the top 10 up-regulated (left) and down-regulated (right) genes in the differential expression analysis. (F) Heatmap showing Pearson correlation values between samples. Samples are clustered according to the grouping (budesonide-treated VS control samples).

**Table S1.** The table shows nuclei count and adipogenic differentiation of *mdx* FAPs for the 8 GCs of the Prestwick library displaying antiadipogenic activity. Values are expressed as percentage compared to DMSO treated cells.

**Table S2.** The table shows the list of significantly modulated genes in *mdx* FAPs treated with 5 μM budesonide for 24 hours (separate excel file).

**Fig. S1**

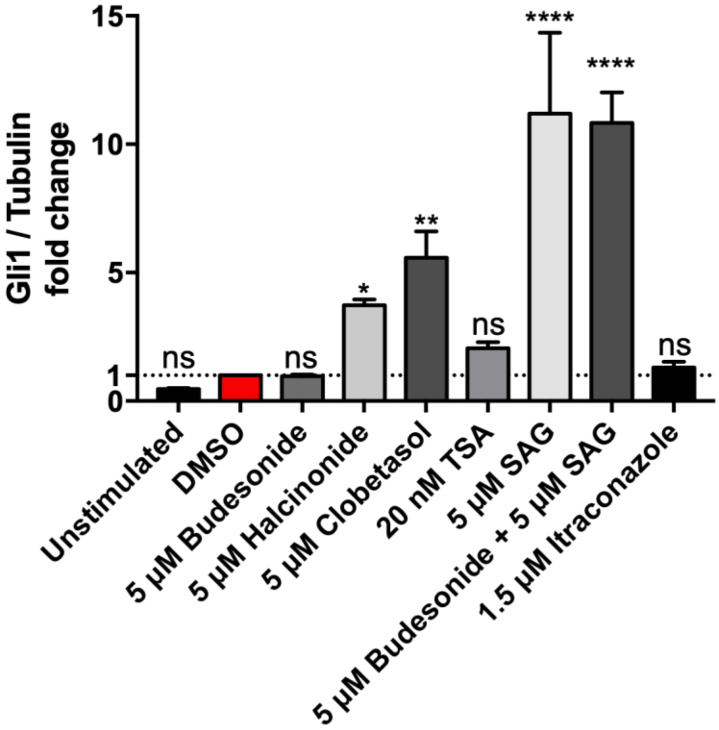

Gli1 / Tubulin fold change
15
10
5
1
0
ns
ns
*
**
ns
****
****
ns
Unstimulated
DMSO
5 µM Budesonide
5 µM Halcinonide
5 µM Clobetasol
20 nM TSA
5 µM SAG
5 µM Budesonide + 5 µM SAG
1.5 µM Itraconazole

Fig. S2

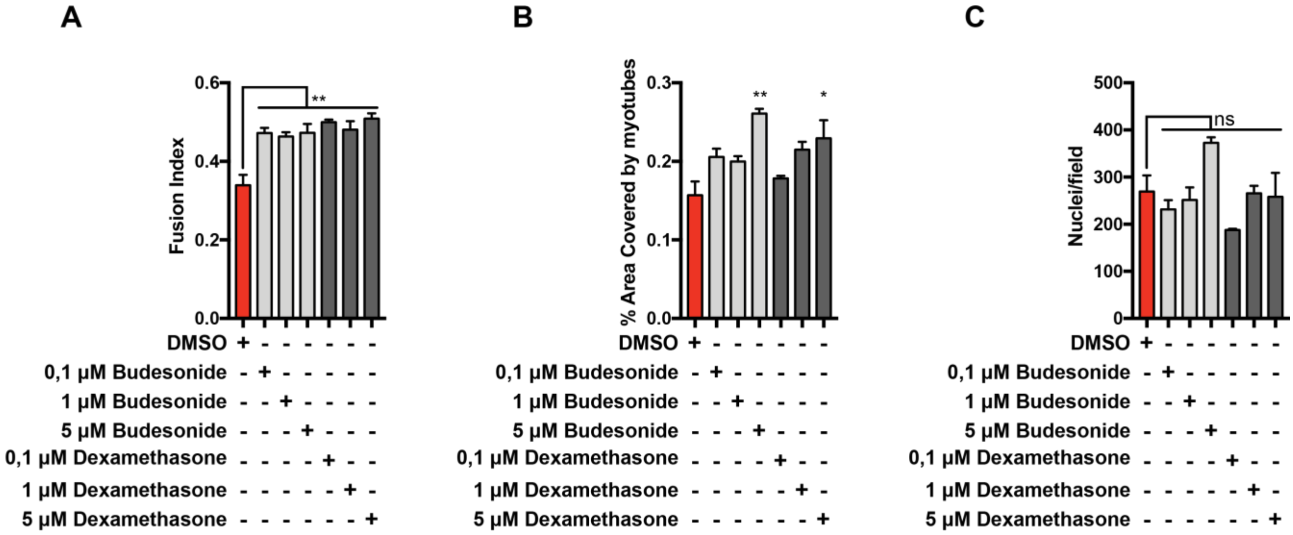
A
B
C
Fusion Index
% Area Covered by myotubes
Nuclei/field
**
*
ns
DMSO
0,1 µM Budesonide
1 µM Budesonide
5 µM Budesonide
0,1 µM Dexamethasone
1 µM Dexamethasone
5 µM Dexamethasone

Fig. S3

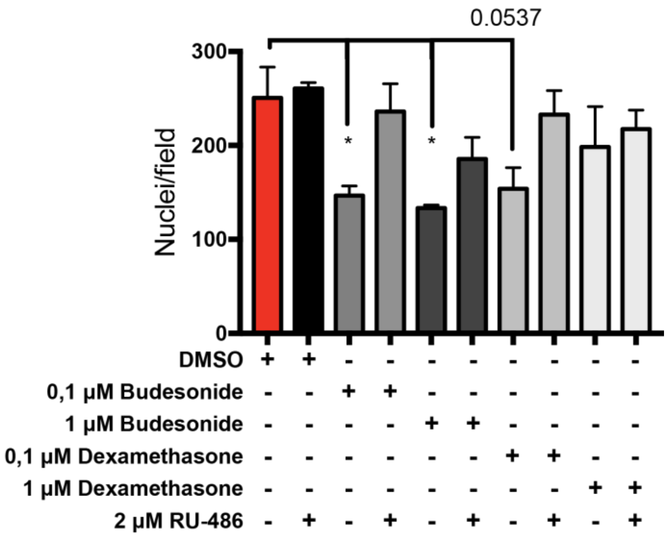
0.0537
Nuclei/field
*
DMSO
0,1 µM Budesonide
1 µM Budesonide
0,1 µM Dexamethasone
1 µM Dexamethasone
2 µM RU-486

**Fig. S4**

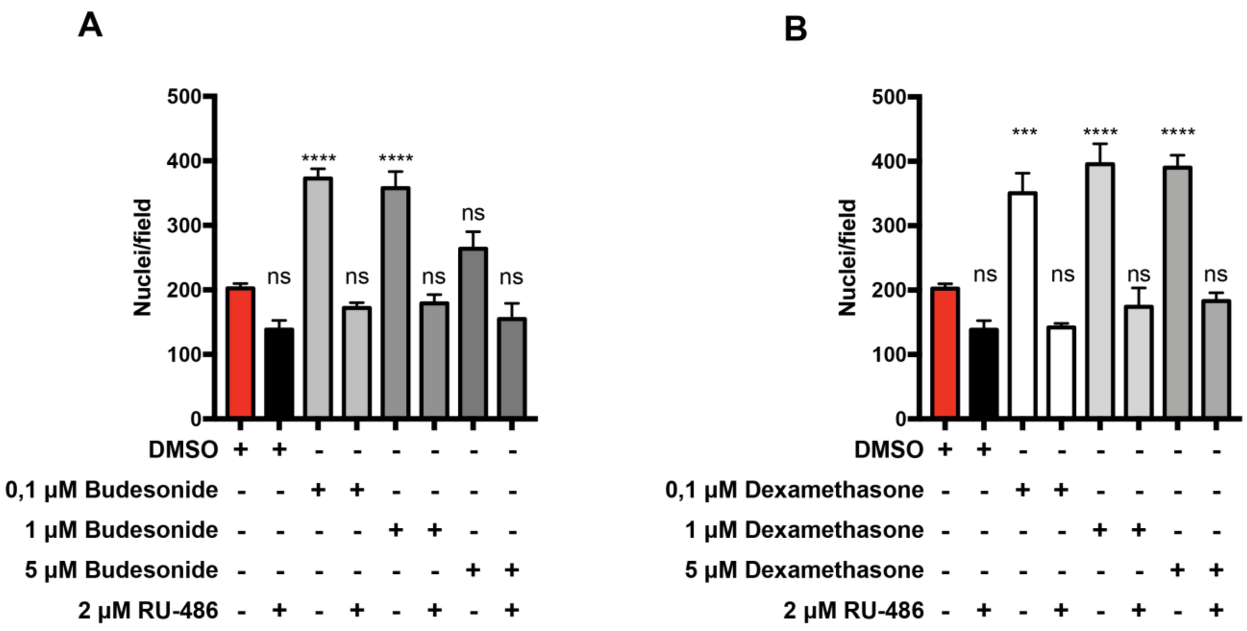
A
Nuclei/field
500
400
300
200
100
0
****
****
ns
DMSO + + - - - - - -
0,1 µM Budesonide - - + + - - - -
1 µM Budesonide - - - - + + - -
5 µM Budesonide - - - - - - + +
2 µM RU-486 - + - + - + - +
B
Nuclei/field
***
****
ns
DMSO + + - - - - - -
0,1 µM Dexamethasone - - + + - - - -
1 µM Dexamethasone - - - - + + - -
5 µM Dexamethasone - - - - - - + +
2 µM RU-486 - + - + - + - +

**Fig. S5**

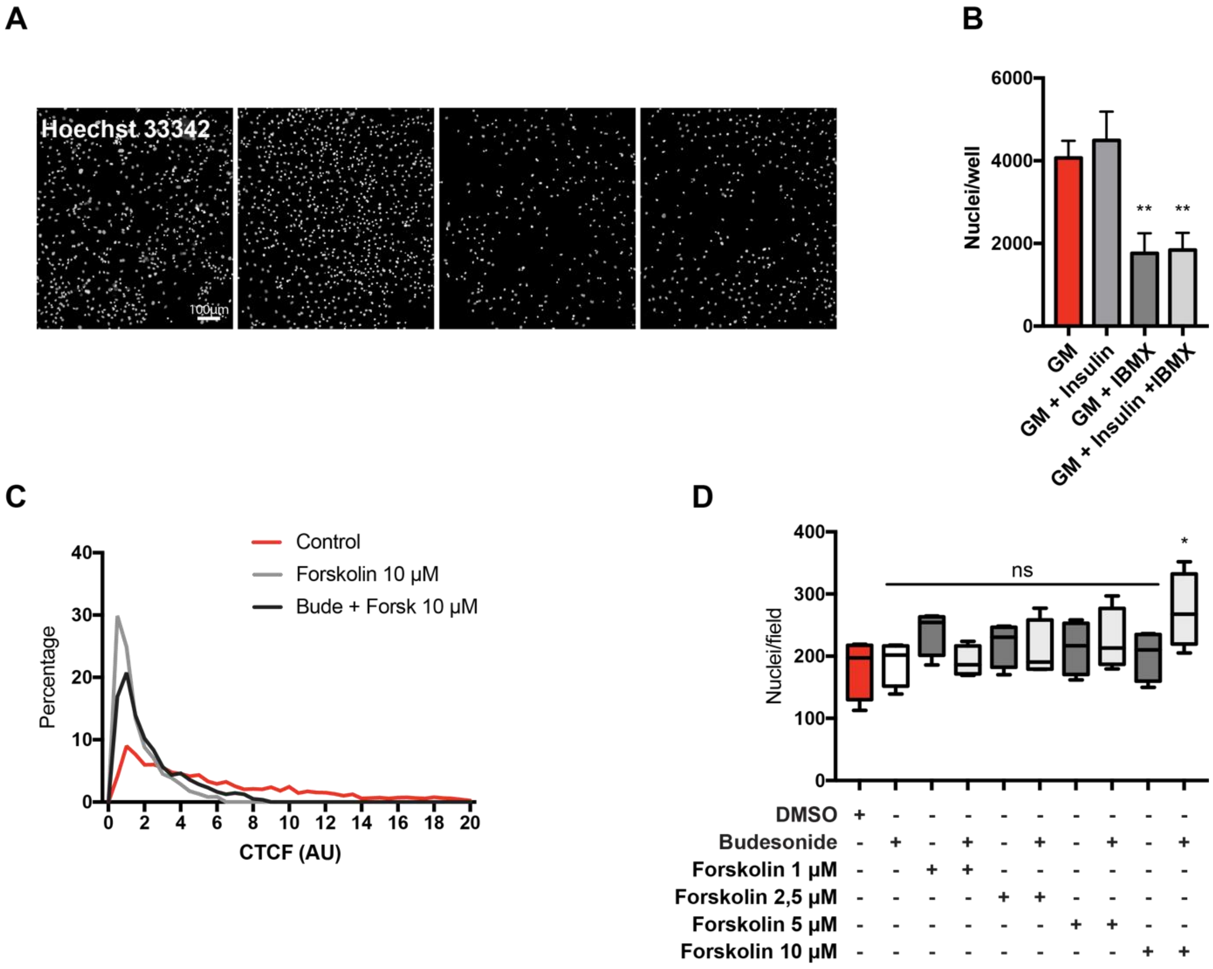
A
Hoechst 33342
100µm
B
Nuclei/well
6000
4000
2000
0
**
GM
GM + Insulin
GM + IBMX
GM + Insulin +IBMX
C
Control
Forskolin 10 µM
Bude + Forsk 10 µM
Percentage
CTCF (AU)
D
Nuclei/field
ns
*
DMSO + - - - - - - - - -
Budesonide - + - + - + - + - +
Forskolin 1 µM - - + + - - - - - -
Forskolin 2,5 µM - - - - + + - - - -
Forskolin 5 µM - - - - - - + + - -
Forskolin 10 µM - - - - - - - - + +

**Fig. S6**

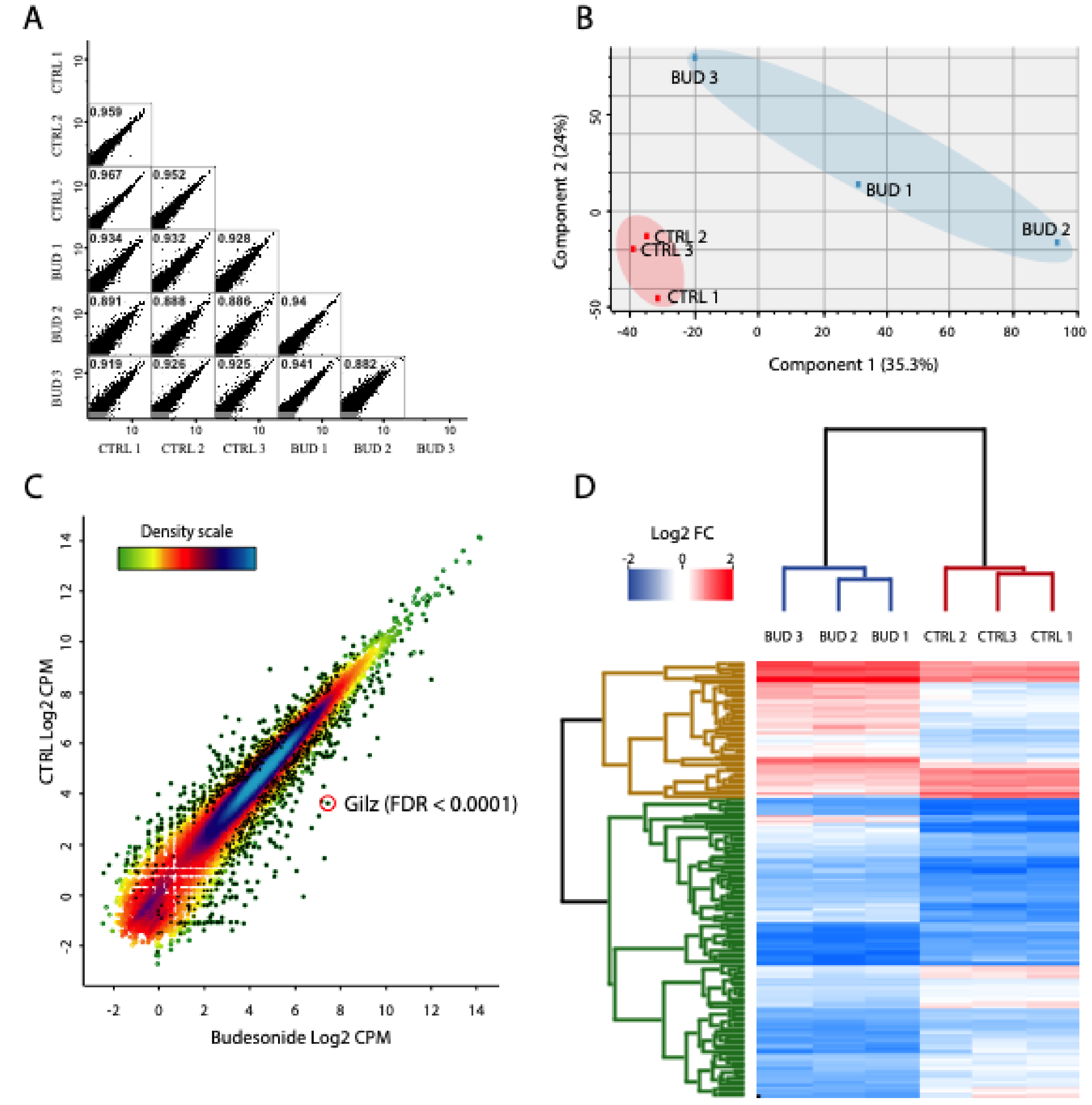


E

| Top 10 up-regulated | Top 10 down-regulated |
|---|---|
| Aldh1a1 | Slc41a2 |
| Map7d2 | Trib2 |
| Fmo2 | Vangl2 |
| Ctla2a | Apol6 |
| Tsc22d3 (GILZ) | Plxdc2 |
| Hsbp7 | Maged1 |
| Ramp2 | Ankrd50 |
| Ndrg2 | Mmp2 |
| Wnk4 | Spry2 |
| Eva1c | Tnk2 |

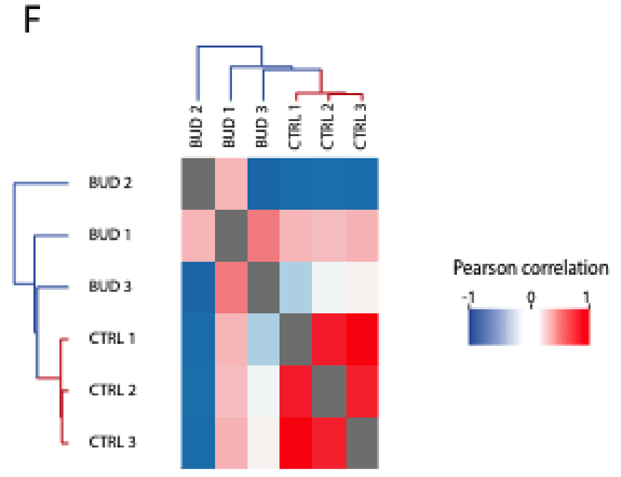

**Table S1**

| | Nuclei count | Adipogenic differentiation |
|---|---|---|
| **Drug Name** | % compared to DMSO | % compared to DMSO |
| MOMETASONE FUROATE | *109,8* | *18,0* |
| HYDROCORTISONE BASE | *115,1* | *37,9* |
| DIFLORASONE DIACETATE | *115,1* | *30,2* |
| FLUMETHASONE | *116,4* | *47,9* |
| HALCINONIDE | *96,4* | *51,6* |
| CLOBETASOL PROPIONATE | *73,2* | *24,3* |
| BUDESONIDE | *89,9* | *1,2* |
| FLUOCINONIDE | 88,5 | 48,5 |
